# Large-scale dimensional behavioral profiling dissociates fear memory from locomotor confounds in mice: The necessity of baseline-normalized metrics

**DOI:** 10.64898/2026.07.12.738100

**Authors:** Daiki X. Sato, Markos M. Chatzigiannis, Hirotaka Shoji, Giovanni Sala, Satoko Hattori, Keizo Takao, Tatsuo Kinashi, Tatsuya Kishino, Sumiyo Morita, Izuho Hatada, Yo Shinoda, Kotaro Hattori, Takeshi Yagi, Akinobu Matsumoto, Hideo Egawa, Shoko Nishihara, Kimiko Shimizu, Koji Ikegami, Maki K. Yamada, Hiroshi Ageta, Mitsutoshi Setou, Hirotaka Tao, Naoto Ueno, Pradeep Bhandari, Ryuichi Shigemoto, Shuji Wakatsuki, Toshiyuki Araki, Akihiro Yamanaka, Hideyuki Mukai, Tadahiro Nagaoka, Masashi Kishi, Shigeki Furuya, Tomomi Yamamoto, Yoshihiro Kubo, Yuichi Iida, Yasuhiro Kazuki, Hideki Enomoto, Motoaki Fukasawa, Nobuteru Usuda, Satoshi Inoue, Kaoru Inokuchi, Tsuyoshi Hattori, Mariko Taniguchi-Ikeda, Tatsushi Toda, Akiko Kubo, Katsuhiro Kawaai, Katsuhiko Mikoshiba, Louie N. van de Lagemaat, Noboru H. Komiyama, Seth G. N. Grant, Tsuyoshi Miyakawa

**Affiliations:** Institute of Life and Environmental Sciences, University of Tsukuba, Ibaraki, Japan; Institute for Advanced Academic Research, Chiba University, Chiba, Japan; Graduate School of Science, Chiba University, Chiba, Japan; Division of Systems Medical Science, Center for Medical Science, Fujita Health University, Aichi, Japan; Department of Psychology, Institute of Population Health, University of Liverpool, Liverpool, UK; Department of Behavioral Physiology, Graduate School of Innovative Life Science, University of Toyama, Toyama, Japan; Department of Behavioral Physiology, Faculty of Medicine, University of Toyama, Toyama, Japan; Institute of Biomedical Science, Kansai Medical University, Osaka, Japan; Division of Functional Genomics, Graduate School of Biomedical Sciences, Nagasaki University, Nagasaki, Japan; Laboratory of Genome Science, Biosignal Genome Resource Center, Institute for Molecular and Cellular Regulation, Gunma University, Gunma, Japan; Laboratory of Environmental Health, School of Pharmacy, Tokyo University of Pharmacy and Life Sciences, Tokyo, Japan; Department of Biobanking, Medical Genome Center, National Center of Neurology and Psychiatry, Tokyo, Japan; Department of Frontier Biosciences, Osaka University Graduate School of Frontier Biosciences, Osaka, Japan; Division of Biological Science, Graduate School of Science, Nagoya University, Aichi, Japan; Department of Biosciences, Graduate School of Science and Engineering, Soka University, Tokyo, Japan; Glycan and Life Systems Integration Center (GaLSIC), Soka University, Tokyo, Japan; Organization for International Education and Exchange, University of Toyama, Toyama, Japan; Department of Anatomy and Developmental Biology, Graduate School of Biomedical and Health Sciences, Hiroshima University, Hiroshima, Japan; Pharmaceutical Sciences at Kagawa, Tokushima Bunri University, Kagawa, Japan; Division for Therapies Against Intractable Diseases, Center for Medical Science, Fujita Health University, Aichi, Japan; Department of Cellular and Molecular Anatomy, Hamamatsu University School of Medicine, Shizuoka, Japan; Institute of Photonics Medicine, Hamamatsu University School of Medicine, Shizuoka, Japan; University of Occupational and Environmental Health, Fukuoka, Japan; National Institutes of Natural Sciences, Tokyo, Japan; Institute of Science and Technology Austria (ISTA), Vienna, Austria; Department of Peripheral Nervous System Research, National Institute of Neuroscience, National Center of Neurology and Psychiatry, Tokyo, Japan; Chinese Institute for Brain Research (CIBR), Beijing, China; Department of Laboratory Medicine, Medical Research Institute Kitano Hospital, Osaka, Japan; Biosignal Research Center, Kobe University, Hyogo, Japan; Division for Therapies against Intractable Diseases, Center for Medical Science, Fujita Health University, Aichi, Japan; Division of Biological Science, Graduate School of Science and Technology, Nara Institute of Science and Technology, Nara, Japan; Department of Bioscience and Biotechnology, Graduate School of Bioresource and Bioenvironmental Sciences, Kyushu University, Fukuoka, Japan; Division of Biophysics and Neurobiology, National Institute for Physiological Sciences, Aichi, Japan; Chromosome Engineering Research Center, Tottori University, Tottori, Japan; Graduate School of Medicine, Kobe University, Hyogo, Japan; School of Medicine, Faculty of Medicine, Fujita Health University, Aichi, Japan; Department of Systems Aging Science and Medicine, Tokyo Metropolitan Institute for Geriatrics and Gerontology, Tokyo, Japan; Department of Biochemistry, Graduate School of Medicine and Pharmaceutical Sciences, University of Toyama, Toyama, Japan; Department of Anatomy and Neuroscience, Nara Medical University, Nara, Japan; Department of Pediatrics, Kochi Medical School, Kochi University, Kochi, Japan; National Center of Neurology and Psychiatry, Tokyo, Japan; Division of Dermatology, Department of Internal Related, Kobe University, Hyogo, Japan; Laboratory of Cell and Tissue Biology, Keio University School of Medicine, Tokyo, Japan; Shanghai Institute for Advanced Immunochemical Studies (SIAIS), ShanghaiTech University, Shanghai, China; Centre for Haemato-Oncology, Barts Cancer Institute, Queen Mary University of London, London, UK; Centre for Regenerative Medicine, University of Edinburgh, Edinburgh, UK; Genes to Cognition Program, Institute for Neuroscience and Cardiovascular Research, University of Edinburgh, Edinburgh EH16 4SB, UK; Simons Initiative for the Developing Brain (SIDB), Institute for Neuroscience and Cardiovascular Research, University of Edinburgh, Edinburgh EH8 9XD, UK; The Patrick Wild Centre for Research into Autism, Fragile X Syndrome & Intellectual Disabilities, Institute for Neuroscience and Cardiovascular Research, University of Edinburgh, Edinburgh EH8 9XD, UK; Muir Maxwell Epilepsy Centre, Institute for Neuroscience and Cardiovascular Research, University of Edinburgh, Edinburgh EH8 9XD, UK

**Author notes:** Corresponding author: Tsuyoshi Miyakawa.

**Keywords:** mouse, fear conditioning, behavioral test, meta-analysis

## Abstract

Fear conditioning is widely used to assess associative memory in mice, yet percent freezing conflates memory with baseline locomotor and anxiety-related traits. A systematic survey of recent studies (2020–2025) found that fewer than 1% statistically integrate locomotor activity into freezing analyses. Here, we address this gap using a large-scale dataset of >10,000 mice across >160 comparisons, including genetic mutations, pharmacological interventions and aging, tested in 15 standardized behavioral paradigms. Conventional freezing scores covaried strongly with general locomotor activity, obscuring memory-related phenotypes. Multiple factor analysis identified two principal behavioral dimensions, locomotor activity and learning/memory: conventional freezing aligned with the locomotor dimension, whereas freezing subtraction and the activity suppression ratio mapped onto the memory dimension and improved detection of synaptic plasticity phenotypes. These analyses show that baseline locomotor normalization is essential for interpreting fear conditioning as a memory assay and provide an open framework for selecting and reporting locomotor-normalized metrics.

## Introduction

Fear conditioning (FC) is one of the most established paradigms in behavioral neuroscience for studying the mechanisms of associative learning and memory ^1–3^. In this paradigm, rodents are exposed to a conditioned stimulus (CS), such as a tone, paired with an unconditioned stimulus (US), such as a mild foot shock. After this pairing, presentation of the CS alone evokes a fear response, typically quantified as freezing, which reflects the suppression of movement during perceived threat. Owing to its simplicity, ease of quantification, and high reproducibility, freezing has been widely adopted as the principal behavioral index of fear memory ^4^.

Despite its widespread use, freezing is influenced by baseline locomotor activity, motor coordination, and anxiety-related traits, which differ markedly among mouse strains and genetic backgrounds ^5^. Consequently, differences in freezing levels may reflect pre-existing behavioral tendencies rather than differences in learning or memory per se, leading to systematic over-or underestimation of fear memory strength. In addition to baseline behavioral variation, freezing is highly sensitive to experimental parameters such as CS–US interval, shock intensity, and training structure ^6,7^. These procedural factors modulate overall immobility levels and interact with strain-dependent activity profiles, further complicating comparisons across experiments and genotypes. Together, these considerations indicate that freezing duration alone conflates memory-related changes with motor and emotional components, limiting its interpretability as a memory-specific index.

Several studies have questioned whether freezing reliably reflects memory strength, noting that immobility can arise from anxiety, altered sensory processing, or motor inhibition independent of associative learning ^8–11^. To address this issue, various approaches have been proposed to account for baseline activity, including normalization procedures and suppression-based indices. In parallel, many studies have included simple checks of baseline activity or shock reactivity and noted that these measures did not differ significantly between groups, before interpreting percent freezing as a memory index ^7,12^. While such checks help rule out gross motor confounds, they do not provide a quantitative adjustment of freezing for continuous, individual-level variation in activity. For example, suppression ratios or related activity-normalized measures have been employed in studies of synaptic plasticity mutants, including CaMKII-related models ^13,14^, and in genetic studies of learning and memory phenotypes that emphasize the importance of interpreting behavioral outcomes in the context of motor and activity-related traits ^15^.

Large-scale behavioral phenotyping efforts have now made it possible to analyze multiple mouse strains under highly standardized conditions across a wide range of behavioral domains ^12,16–21^. Within this framework, fear conditioning should not be viewed as a simple readout of memory, but as a component of a multidimensional behavioral architecture that includes locomotion and emotional reactivity. This perspective is particularly relevant given the widespread use of fear conditioning in behavioral screening of mouse models for neuropsychiatric and neurodegenerative disorders, where behavioral metrics are expected to be both sensitive and specific to learning and memory processes.

In the present study, we re-evaluated conventional freezing-based metrics using a large-scale behavioral dataset comprising 166 mouse groups, which included genetic mutants, drug-administered, psycho-stimulated, or in aged conditions, tested across 15 standardized paradigms. We first quantified the extent to which conventional freezing scores covary with general locomotor and anxiety-related traits. We then validated baseline-normalized indices, including freezing subtraction and activity suppression ratios, as tools for isolating memory-related variance. Finally, we applied multiple factor analysis (MFA) to position fear conditioning measures within the broader behavioral structure. Through this empirically grounded, dimensional approach, we aim to provide a robust framework for disentangling learning-related signals from locomotor confounds in fear conditioning, thereby enhancing the interpretability and reproducibility of large-scale behavioral phenotyping.

## Results

### Conventional freezing indices covary with locomotor activity across strains

To evaluate the reliability of freezing as a measure of fear memory, we analyzed a large-scale behavioral dataset encompassing 166 mouse groups, including genetic mutants and mice subjected to drug treatments, psychostimulant administration, or aging (see Tables S1 and S2). For each strain, behavioral phenotypes were assessed across 15 standardized paradigms (see Materials and methods), including fear conditioning (FC), elevated plus maze (EP), open field (OF), light–dark box (LD), social interaction tasks (SI), and cognitive assays such as the Barnes maze (BM) and T-maze (TM). Behavioral effects were quantified as Hedges’ *g* between mutant and control littermates, allowing standardized comparisons across assays and strains. We refer to this comprehensive dataset as the mouse behavioral phenotype dataset (MBPD).

An overview of the FC procedure and representative time courses of freezing during training and test sessions for control and *Atf5* knockout (*Atf5*-KO) mice are shown in Fig. 1a. These data illustrate that apparent genotype differences in freezing can emerge at multiple time points, motivating a systematic evaluation of whether such differences reflect memory processes or baseline behavioral traits. We first examined correlations between FC freezing measures and indices of general locomotor activity derived from other behavioral tests, using strain-level effect sizes. Across the MBPD, conventional freezing metrics were significantly associated with multiple activity-related variables across assays (Fig. S1a). Compared to representative locomotor measures (traveled distance in OF and HC; Fig. S1b), EP total distance showed the most consistent negative association with both contextual and cued freezing on Day 2 and was therefore selected for display in Fig. 1b. Strains with higher activity tended to show lower freezing, a relationship that may overlap with learning-related variation but is not readily attributable to memory performance alone. Given that EP performance reflects exploratory drive and anxiety-related avoidance, we next tested whether freezing–activity covariation was strongest during the initial exploratory phase of locomotor behavior by analyzing OF distance in 5-min bins. Correlations with freezing (and with EP activity) were strongest early in the OF session and weakened over time (Fig. S1c), suggesting that early exploration and/or anxiety-related inhibition contributes to the shared variance between conventional freezing measures and locomotor traits. Similar patterns were observed for freezing measured on Day 30, indicating that the association between conventional freezing and activity-related phenotypes persists across longer retention intervals.

**Figure 1.**
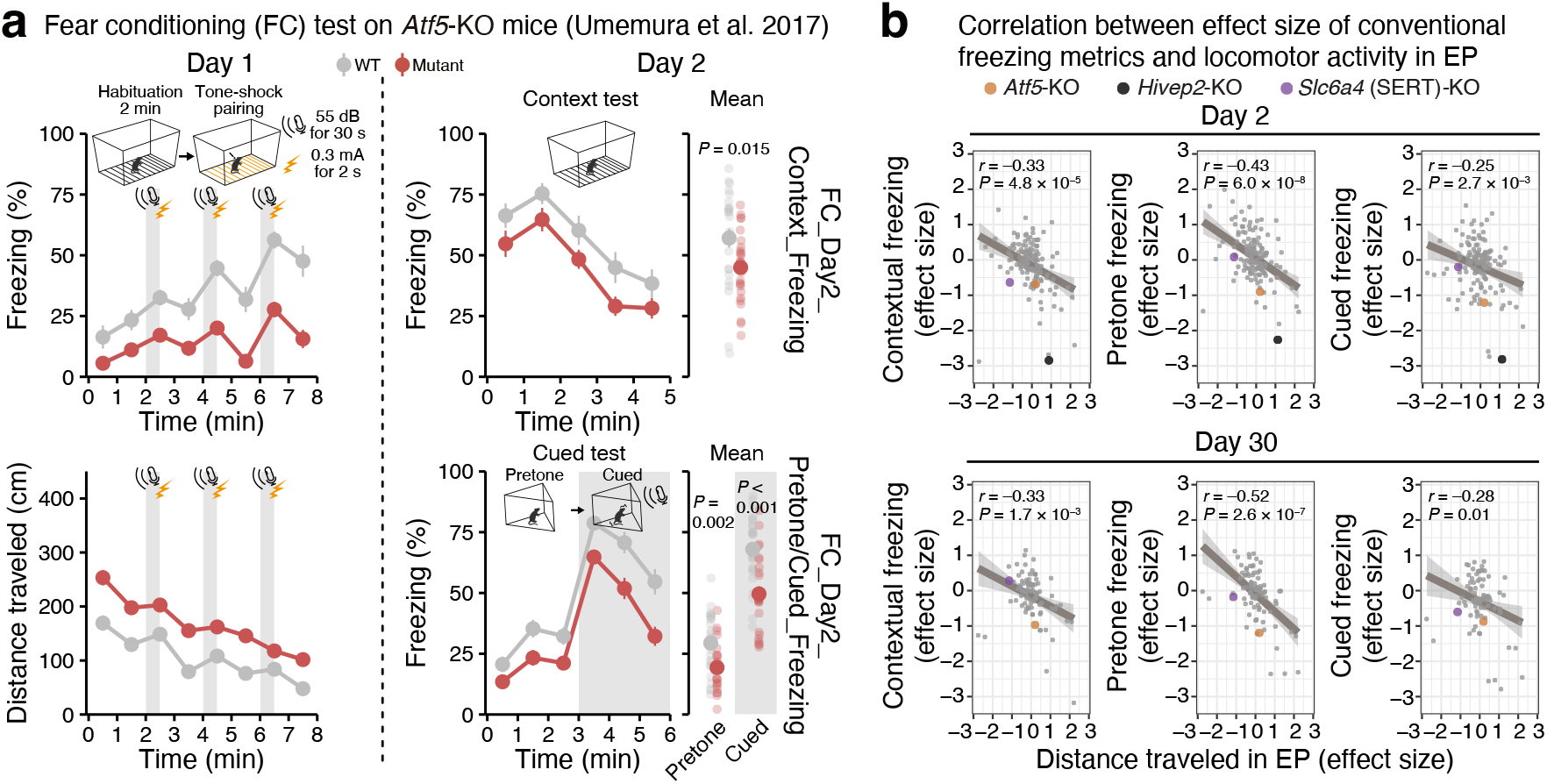
Conventional freezing measures in fear conditioning are confounded by baseline locomotor activity. **(a)** Schematic of the fear conditioning test protocol and calculation of conventional freezing metrics. On Day 1, mice underwent a habituation period followed by tone-shock pairings. On Day 2 or Day 30, context and cued tests were performed. Example fear conditioning data from the *Atf5*-KO mutant line are shown ^29^. On Day 1, freezing responses increased during tone–shock pairings (top left), while distance traveled exhibited sharp increase in response to the shock (bottom left, mean of three trials). On Day 2, context and cued (but in pre-tone phase) tests revealed differences in freezing. These data illustrate the derivation of conventional immobility-based metrics (i.e., context, pretone, or cued freezing). **(b)** Correlations between conventional freezing metrics on Days 2 and 30 and total distance traveled in the elevated plus maze (EP_TotalDistance). Across strains, freezing scores were consistently and significantly negatively correlated with general activity, highlighting the confounding influence of locomotor traits on fear memory assessments.

### Baseline-normalized metrics minimize activity dependence and increase sensitivity

We next re-evaluated FC performance using baseline-normalized metrics designed to reduce confounding by strain-dependent differences in general activity. Two indices were calculated: (1) Freezing Subtraction, defined as the difference in freezing between the test and training sessions, which is referred to as “memory mean” in a previous study ^19^, and (2) Activity Suppression Ratio, defined as the relative reduction in movement during the test compared with baseline activity ^14^ (Fig. 2a). While conventional freezing scores revealed significant differences between *Atf5*-KO and control mice (Fig. 1a), these differences were not observed when either baseline-normalized metric was applied (Fig. 2b). Moreover, both normalized indices showed substantially weaker correlations with locomotor activity than did conventional freezing measures (Fig. 2c). At the individual mouse level, conventional freezing measures remained negatively associated with activity indices, although the effect sizes of these correlations were modest rather than strong (Fig. S2a). In contrast, both baseline-normalized indices exhibited minimal dependence on locomotor activity (Fig. S2b). These results indicate that the confounding effect of locomotion is evident at both the strain and individual levels but can be attenuated by normalization procedures.

**Figure 2.**
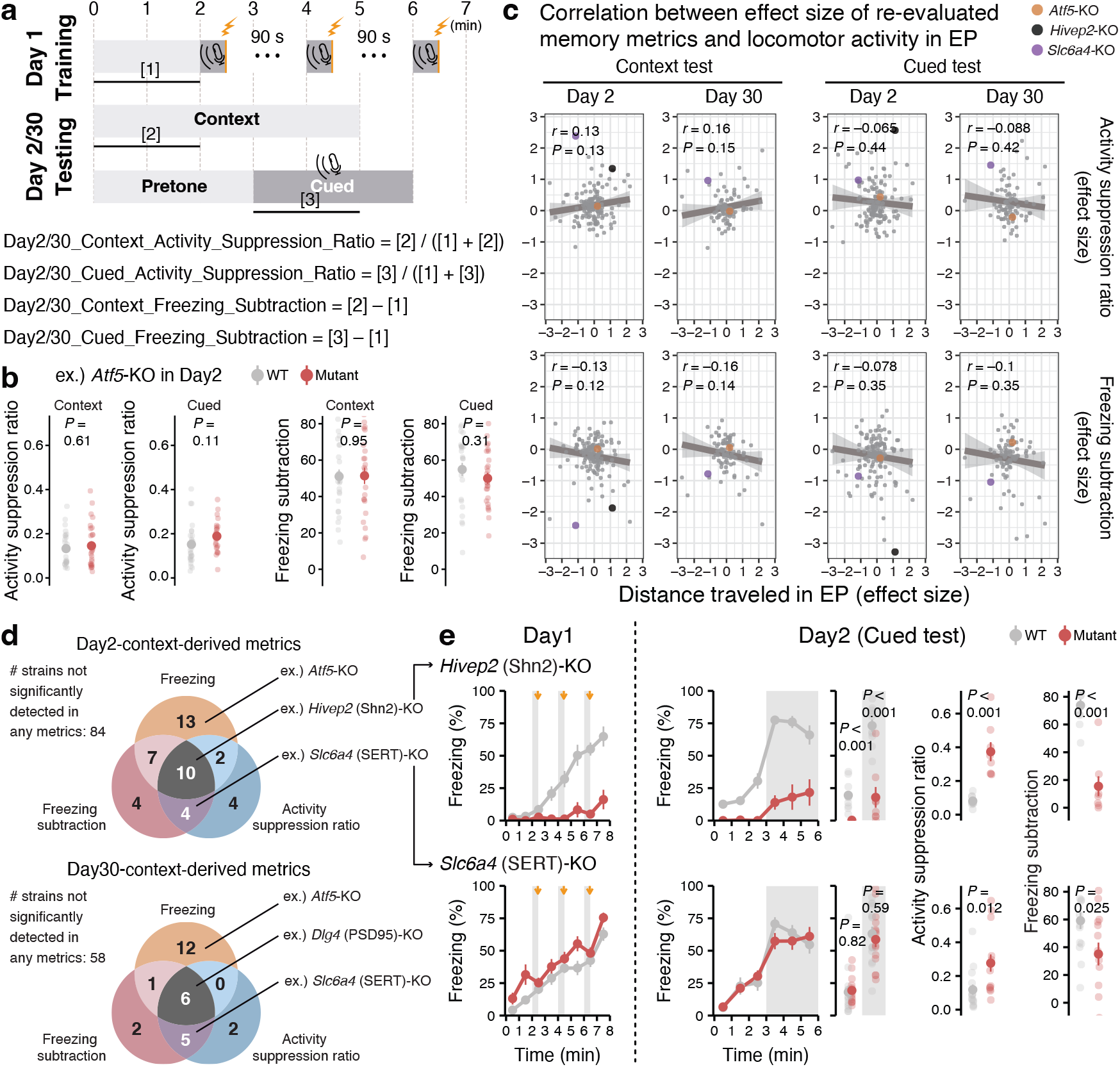
Baseline-normalized fear-conditioning metrics reduce locomotor confounds and enhance detection of memory-related phenotypes. **(a)** Re-evaluated freezing metrics were calculated either as the ratio of test-period traveled distance to the total distance (test + baseline) (Activity Suppression Ratio ^14^) or the difference (subtraction) in freezing levels between baseline and during training session (Freezing Subtraction ^19^). **(b)** The baseline-normalized metrics showed no significant difference between control and mutant lines of *Atf5*-KO and **(c)** no significant correlation with general locomotor activity, as measured by total distance traveled in the elevated plus maze (EP_TotalDistance). **(d)** Venn diagrams show the number of strains with significant behavioral differences between control and experimentally treated groups for context tests on Day 2 (top) and Day 30 (bottom), as assessed using three different metrics: conventional freezing scores, freezing subtraction scores, and activity suppression ratios. Notably, the baseline-normalized metrics detected unique sets of strains and showed greater overlap with each other compared to the conventional freezing metric. Representative strains in each overlap region are listed. **(e)** Representative examples of cued-test phenotypes detected using conventional and baseline-normalized metrics. Freezing time courses during Day 1 training and Day 2 cued testing are shown for *Hivep2* (Shn2)-KO and *Slc6a4* (SERT)-KO mice. Gray shaded periods indicate cue presentations, and lightning symbols indicate foot shocks during training. Summary plots compare conventional freezing scores, activity suppression ratios, and freezing subtraction scores between WT and mutant mice in the Day 2 cued test. *Hivep2*-KO showed significant differences across all three metrics, whereas *Slc6a4*-KO showed no significant difference in conventional freezing but was detected by the baseline-normalized metrics, illustrating how these measures can reveal memory-related phenotypes that are obscured in raw freezing scores.

To assess how metric choice influences the detection of genotype effects, we systematically compared significant genotype effects across all combinations of FC metrics and test conditions (context vs. cued; Day 2 vs. Day 30). Figures 2d and S3 summarize these comparisons, with Venn diagrams quantifying overlap across metrics (Fig. 2d) and strain-level annotations identifying the genotypes populating each region across conditions (Fig. S3). Multiple well-established mutants of the genes involved in synaptic plasticity, including *Grin1*, *Grin2a*, *Camk2a*, *Dlg4*, *Gria1*, *Ppp3r1*, and *Hivep2*, all of which are well known to impair learning and memory ^22–28^, were detected robustly across both conventional and baseline-normalized metrics (colored in grey), indicating their strong phenotypes that persist even after baseline correction. Notably, *Slc6a4* (SERT)-KO was detected only by the baseline-normalized metrics (colored in purple) (Figs. 2d–e). Because these mice are hypoactive at baseline, conventional freezing measures can mask memory-relevant genotype effects. By accounting for baseline activity, the normalized metrics reduce this risk of false negatives in genetic screens. Conversely, several strains, including *Atf5*-KO, were significant only with conventional freezing measures and lost significance when baseline-normalized indices were applied (colored in orange), suggesting that their apparent impairments may be driven primarily by altered activity or anxiety-related traits rather than memory-specific effects ^29–31^. These analyses demonstrate that baseline normalization improves specificity for memory-related phenotypes while reducing potential false positives due to general activity differences.

### Multiple factor analysis identifies two principal behavioral dimensions

To contextualize FC metrics within the broader structure of behavioral variation, we conducted multiple factor analysis (MFA) across 15 standardized tests comprising 68 behavioral variables (Fig. 3a). The first two dimensions accounted for 33.51% of the total variance (20.14% for Dimension 1 and 13.37% for Dimension 2; Fig. 3b). Although the dataset included diverse behavioral readouts, a substantial fraction of variance was captured by this low-dimensional structure. Dimension 1 primarily captured general locomotor activity and anxiety-like behavior, with high loadings for measures such as total distance traveled in OF, EP, CSI, and SI. Dimension 2 represented learning and memory-related performance, integrating measures from FC, BM, TM, and Porsolt forced swim test (PS) (Fig. 3b). Thus, the two dominant dimensions separated activity–anxiety traits from cognitive performance.

**Figure 3.**
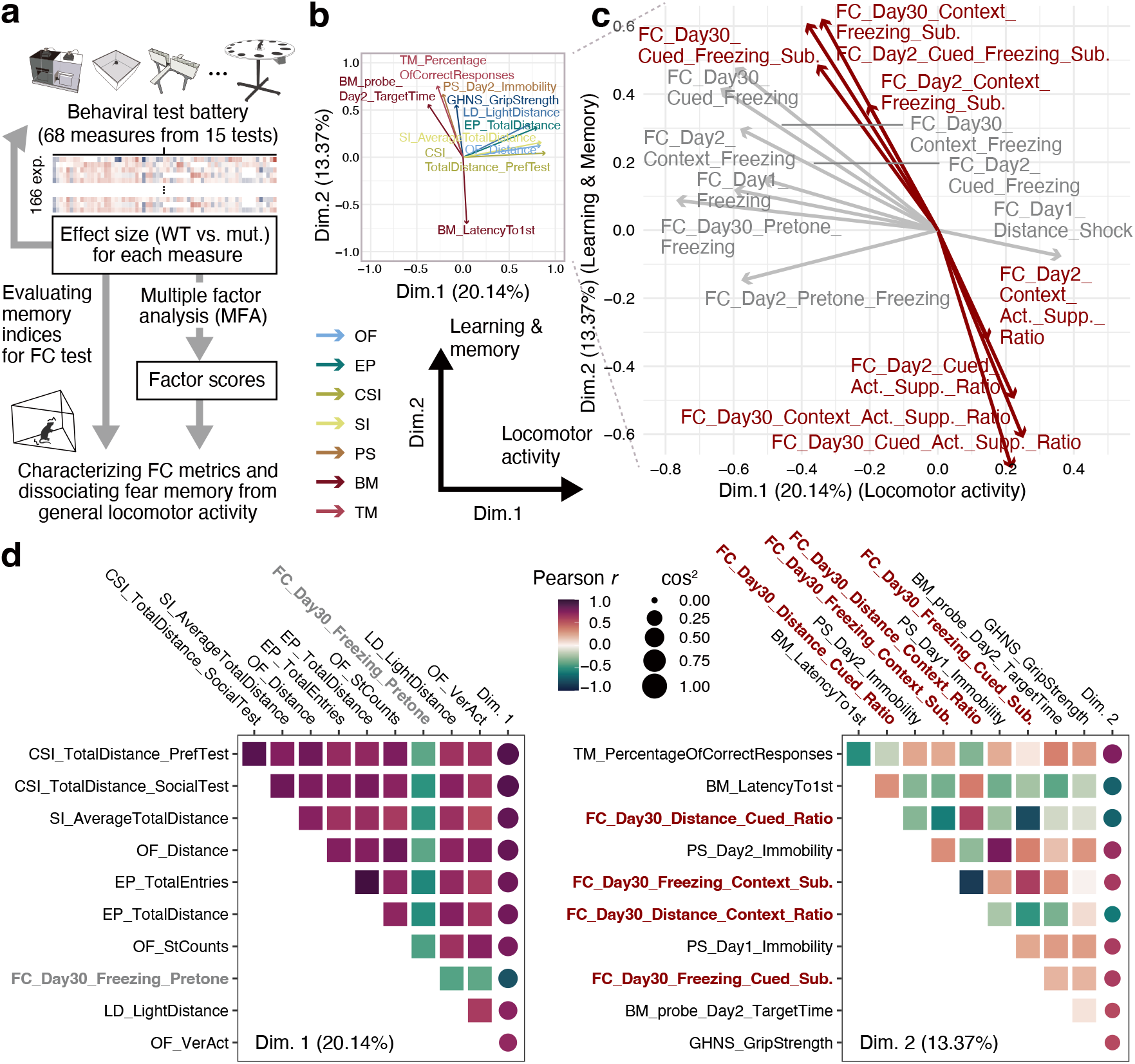
Multiple factor analysis reveals a two-dimensional organization of mouse behavior separating locomotor and cognitive domains. **(a)** Overview of the analytical framework. Behavioral data were collected from 166 mouse groups, which included genetic mutants, drug-administered, psycho-stimulated, or in aged conditions. Across 15 tests, 68 behavioral measures were yielded, which were used to calculate effect sizes (Hedges’ *g*) for mutant versus wild-type (WT) comparisons. After the imputation of missing values, multiple factor analysis (MFA) was used to derive latent behavioral dimensions. **(b)** Results of a multiple factor analysis (MFA) show that the first dimension (Dim. 1) primarily captures locomotor activity, while the second dimension (Dim. 2) reflects memory and cognitive performance. **(c)** Projection of only the freezing-related variables from the MFA reveals that conventional freezing metrics cluster along the locomotor dimension (Dim. 1), whereas the proposed metrics align more strongly with the cognitive/memory dimension (Dim. 2). **(d)** Correlation matrix showing the relationship between behavioral variables and the first two MFA dimensions. The plot highlights that Dim. 1 is strongly associated with locomotor-related variables, while Dim. 2 is associated with memory-related measures such as those from the T-maze (TM) and Barnes maze (BM), as well as the baseline-normalized freezing metrics.

Notably, when focusing on FC-related variables (Fig. 3c), the conventional freezing metrics predominantly extended along Dimension 1, indicating strong alignment with activity-anxiety-related variance. In contrast, baseline-normalized FC indices projected along Dimension 2, consistent with a closer association with learning and memory performance. Quantitative correlations between each behavioral variable and the two dimensions (Fig. 3d) further showed that remote fear memory measures, namely FC indices assessed on Day 30, together with TM and BM indices, ranked among the strongest associations with Dimension 2, whereas Day 2 freezing measures showed weaker associations. This pattern suggests that remote memory performance in FC captures memory-related variance more robustly than near-term measures obtained at 24 hours. Interestingly, the immobility in the PS also showed a strong correlation with Dimension 2. Traditionally interpreted as an index of depressive-like behavior, PS immobility may additionally reflect learning-related processes ^32–34^. One possible explanation is that high immobility early in the test requires learning that the situation is inescapable, thereby engaging memory-related mechanisms rather than purely affective states.

### Cross-validation with independent G2C data confirms behavioral dimensionality

To evaluate the reproducibility of the behavioral dimensions identified in the MBPD dataset, we conducted a cross-validation analysis using an independent behavioral resource, the Genes-to-Cognition (G2C) database ^16^. The G2C project systematically generated knockout mice for synapse-related genes and subjected them to a standardized battery of behavioral assays, providing an ideal external benchmark. Both datasets included overlapping variables from OF, EP, RR and FC, which allowed direct comparison of multivariate structures. Notably, the G2C fear-conditioning measures correspond to baseline-normalized or activity-controlled indices ^19^, facilitating alignment with our refined metrics.

When MFA was performed separately within each dataset using the shared variables (EP_TotalDistance, EP_OpenArmTimePercentage, OF_Distance, RR_Learning, RR_Memory, and FC_Day2_Context_Freezing_Sub.), both analyses revealed a similar two-dimensional configuration (Fig. 4). In both the MBPD and G2C datasets, Dimension 1 primarily reflected locomotor activity, whereas Dimension 2 corresponded to learning and memory performance. The spatial arrangements of key variables including FC subtraction measures and learning indices from RR were preserved across datasets, indicating a consistent behavioral architecture. Notably, EP_OpenArmTimePercentage was positioned closer to Dimension 2 in the MBPD analysis than in G2C (Fig. 4a), suggesting that the anxiety-related component captured by open-arm exploration may partially covary with learning-related variance in the dataset, even though the overall two-axis structure was conserved.

**Figure 4.**
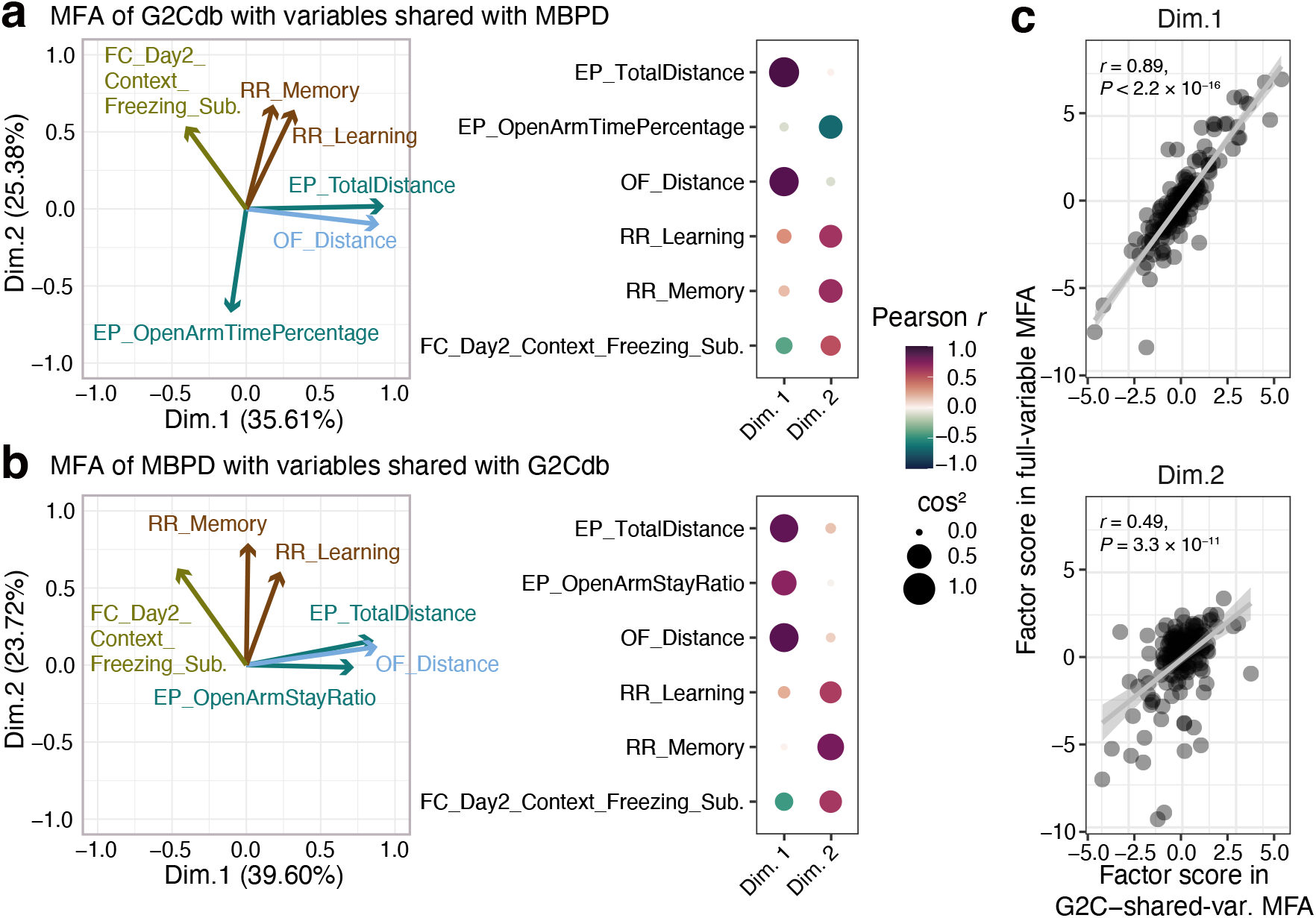
Cross-validation of behavioral dimensions across independent datasets. Multiple factor analysis (MFA) of six shared behavioral variables in **(a)** Genes to Cognition Database (G2Cdb) of 60 strains and **(b)** our dataset of 166 mouse groups. (Left) Dimension 1 primarily captures general locomotor activity, with high loadings from total distance traveled in the elevated plus maze (EP_TotalDistance) and open field (OF_Distance), while Dim. 2 reflects memory-related performance, with contributions from rotarod learning (RR_Learning), rotarod memory (RR_Memory), and the fear conditioning subtraction metric (FC_Day2_Context_Freezing_Sub.). (Right) Correlation matrix showing the relationship between individual variables and the two dimensions. **(c)** Correlation of factor scores between MBPD analyses using all 68 behavioral variables and those restricted to the six variables shared with the G2C database. Each point represents a mouse strain. Both Dimension 1 (locomotor activity; top) and Dimension 2 (learning–memory performance; bottom) showed strong positive correlations (*r* = 0.89 and *r* = 0.49, respectively), indicating that the same two-dimensional behavioral structure is preserved even when a small subset of representative assays is used. This result demonstrates that combinations of a few tasks encompassing locomotor and cognitive components can capture the broader behavioral architecture observed across the full test battery.

Factor scores derived from the MBPD using all 68 variables were highly correlated with those obtained using only the six variables shared with G2C (Fig. 4c). Strong correlations were observed for both Dimension 1 (*r* = 0.89) and Dimension 2 (*r* = 0.49), demonstrating that the core behavioral structure can be recovered even from a reduced variable set. These results also support the robustness of the two-dimensional structure, in which locomotor activity constitutes the primary axis and learning–memory performance forms the secondary axis. Importantly, within this reproducible behavioral architecture, baseline-normalized fear-conditioning metrics consistently aligned with the learning–memory dimension rather than the locomotor dimension. Together, these findings validate the generality of the behavioral framework and provide direct support for the central premise of this study: that refined fear-conditioning metrics offer a more specific and robust representation of memory-related variance than conventional freezing measures, independent of confounding effects of locomotor activity.

### Literature analysis reveals limited integration of locomotor controls in fear conditioning studies

Finally, to evaluate how explicitly locomotor activity is incorporated into the statistical analysis of fear conditioning outcomes, we conducted a systematic literature survey of mouse fear conditioning studies. Automated text-mining of the Europe PMC corpus identified 2803 experimental articles (2020-2025) reporting fear conditioning with accessible full text (see Table S3). Studies were categorized based on how locomotor data were treated in relation to freezing behavior, with manual confirmation for those initially classified as involving explicit statistical integration of locomotor activity (Categories 1 and 2a; see Materials and Methods). Of these, 1932 studies (68.93%) reported performing at least one locomotor or exploratory activity test, most commonly the open field test, elevated plus maze, or light–dark box, while 871 studies (31.07%) did not mention locomotor assessment. Manual review confirmed that only 12 studies (0.43%) explicitly integrated locomotor activity into freezing analyses, including 10 studies using locomotor-adjusted indices and 2 studies incorporating locomotor activity as a statistical covariate in freezing analyses. The remaining 1920 studies (68.50%) reported locomotor measures independently of freezing performance. Together, these results indicate that although locomotor activity is frequently measured in fear conditioning studies, it is rarely incorporated into the statistical evaluation of freezing behavior.

## Discussion

This study re-evaluated fear conditioning (FC) as a measure of memory by integrating behavioral data from more than 160 mouse groups across 15 standardized tasks. By analyzing this large-scale dataset, we revealed that percent freezing, the classical measure of FC, is confounded by baseline locomotor activity and that baseline-normalized indices provide a more reliable representation of learning and memory. The multivariate framework established here clarifies how FC relates to other behavioral domains and highlights locomotor activity and memory performance as two principal axes of behavioral variation. These findings not only refine the interpretation of FC but also contribute to a broader understanding of behavioral organization in mice.

### Reinterpreting freezing as a behavioral composite

Freezing has traditionally been interpreted as a direct measure of associative memory strength ^1,4,8,9^, yet our results demonstrate that this assumption overlooks substantial variance introduced by baseline motor traits. In many conventional studies, locomotor activity has been assessed separately and reported as showing no significant group differences, after which percent freezing is used as a primary memory index ^7,26,35^. While this practice guards against gross locomotor confounds, it does not address continuous, individual-level variation in activity and anxiety that can systematically bias freezing measures. Our findings indicate that quantitative normalization at the level of individual or strain-specific activity, rather than binary significance testing of activity differences, is critical for isolating memory-related variance. In our dataset, conventional freezing scores covaried with multiple locomotor indices, including total distance traveled in EP, OF, CSI, SI (Figs. 1b and S1), consistent with earlier reports that activity differences across strains confound memory assessment ^36^. This suggests that immobility during FC reflects a composite of learned fear, motor output, and anxiety-related inhibition ^10,37,38^.

By applying two baseline-normalized indices, Freezing Subtraction and Activity Suppression Ratio ^14,19^, we minimized these confounds. These measures showed little correlation with locomotor activity (Fig. 2c) and effectively distinguished genotype effects in well-established learning mutants. Some of our previous studies have already adopted activity-normalized measures in specific genetic models, including *Rapgef6*-deficient mice ^31^, underscoring that the conceptual importance of normalization has been recognized. Our contribution lies not only in proposing alternative behavioral indices per se, but in systematically evaluating previously proposed yet underutilized normalization approaches across a large and diverse behavioral space. Across strains, baseline normalization reduced apparent FC deficits in genotypes with prominent locomotor or stress-related alterations, exemplified by *Atf5* and *Reln* mutants ^29,39^, which were primarily detected by conventional freezing metrics alone. In contrast, several canonical learning-related genes implicated in synaptic plasticity and disorders with cognitive dysfunctions, including *Grin1*, *Camk2a*, *Dlg4*, *Gria1*, and *Hivep2*, remained robustly detected after baseline correction ^22–28^. Baseline normalization can also reveal learning-related effects that may be obscured when genotypes differ in baseline activity or stress reactivity. For example, in *Slc6a4* (SERT) and *Ube3a* mutant lines, both of which show prominent non-learning phenotypes that can influence freezing ^40,41^, genotype differences were captured more consistently by baseline-normalized measures than by raw freezing alone (Figs. 2d–e and S3). Collectively, these findings suggest that baseline normalization improves the interpretability of fear conditioning by reducing activity-driven misclassification while preserving, and in some cases enhancing, sensitivity to genuine learning and memory phenotypes, demonstrating their necessity for disentangling memory-related signals from locomotor confounds.

### A robust two-dimensional architecture of mouse behavior

Multiple factor analysis (MFA) of 68 variables across 15 standardized tests revealed two principal behavioral dimensions that together explained approximately one-third of the total variance (Fig. 3). Dimension 1 primarily captured variance related to locomotor and exploratory activity together with anxiety-like behavior, whereas Dimension 2 was predominantly associated with learning-and memory-related performance. Importantly, FC variables, when corrected for baseline activity, mapped onto Dimension 2 alongside BM and TM measures, indicating that these assays share a common cognitive component. Notably, TM is typically interpreted as indexing working memory, whereas other tasks including BM and FC are more closely related to reference memory. Therefore, the fact that both sets of measures map onto Dimension 2 in the present MFA does not necessarily imply that they rely on an identical underlying mechanism. Rather, it suggests that a common latent cognitive component captured by Dimension 2 may be influenced by the same genetic perturbation, such that the mutation can affect both reference-and working-memory–related performance. Interestingly, immobility in PS, a classic measure of depressive-like behavior, also correlated with Dimension 2. This observation suggests that behavioral inhibition in PS may not solely index affective despair but may partially reflect learning-related processes, such as the acquisition of inescapability of the context or adaptive control over behavioral output^32–34^.

Conventional freezing measures without baseline correction were preferentially aligned with Dimension 1. The stronger alignment of fear conditioning measures with Dimension 2 from Day 2 to Day 30 suggests that longer retention intervals may be more sensitive to genetic perturbations that affect cognitive-performance-related processes than 24-hour freezing responses. Notably, in contextual fear conditioning, freezing at ∼24 hours is often treated as an index of recent memory, whereas freezing after longer intervals (e.g., weeks) is used to probe remote memory; these timepoints can differ not only quantitatively but also in the degree to which they rely on partially distinct retrieval and consolidation processes (e.g., systems consolidation ^42,43^). From this perspective, our stronger association of freezing with Dimension 2 at Day 30 is consistent with the idea that remote-memory retrieval more strongly reflects stabilized mnemonic representations, whereas recent-memory freezing may be more susceptible to non-mnemonic contributors such as acute stress reactivity, arousal, and motor inhibition at test ^44,45^. In other words, 24-hour freezing, while widely used as a standard retention measure, may still capture a mixture of mnemonic and immediate performance factors, whereas longer-term recall may be more sensitive to genetic influences on cognitive-performance-related processes.

Importantly, the two-axis organization was not specific to our cohort or analytic choices. We observed the same two-dimensional structure in an independent dataset from the G2C project ^16^, which systematically phenotyped knockout mice for synapse-related genes using a standardized behavioral battery. Within MBPD, factor scores derived from all 68 variables and from only six variables shared with the G2C dataset remained highly correlated (*r* = 0.89 for Dimension 1; *r* = 0.49 for Dimension 2; Fig. 4c). This cross-dataset agreement indicates that the activity-and cognition-related axes capture a robust behavioral scaffold that persists even under substantial reduction of the variable set and across independent large-scale pipelines. Consistent with this view, relatively few prior studies have explicitly modeled animal behavior as orthogonal dimensions of activity and cognition; however, multiple data-driven analyses have independently recovered latent components that closely resemble this structure. Exploratory factor analyses of open-field behavior identified independent components corresponding to overall locomotor output and spatial or exploratory organization, effectively separating activity from other behavioral factors ^46^. Broader behavioral batteries analyzed using multivariate approaches likewise extracted latent components dominated by motor output versus task performance, which together explain genotype-dependent behavioral variation ^47^. Conceptually, our results align with frameworks that organize mouse behavior along continua spanning exploratory drive to higher-order cognitive control ^12,48^. Recent data-driven analyses likewise dissociate movement-related from task-related behavioral structure, with some components tracking engagement or performance and others relating more closely to learning ^49–51^. Collectively, these studies converge with our findings, supporting the view that rodent behavior can be described within a low-dimensional architecture whose primary axes represent locomotor and cognitive domains. Importantly, positioning FC within this manifold clarifies why conventional freezing often yields ambiguous phenotypes and why baseline-normalized FC metrics align more specifically with the cognitive axis.

### Implications for behavioral phenotyping and systems neuroscience

The identification of locomotor and cognitive axes provides a conceptual and quantitative framework for interpreting large-scale behavioral data. Traditional approaches to mouse phenotyping often rely on single-task readouts, which can obscure the underlying structure of behavior and complicate comparisons across studies or laboratories ^12,17^. By contrast, the dimensional framework described here enables behavioral measures to be placed within a shared coordinate space, allowing more precise inference about the processes underlying each task. This approach parallels recent advances in the “big behavior” movement, which emphasizes high-throughput, data-driven quantification and the recovery of latent behavioral structure or phenotypes from rich behavioral data ^52,53^. Integrating such dimensional representations with genetic and neural data may ultimately reveal how molecular and circuit-level variation maps onto the fundamental organizational principles of behavior.

From a methodological perspective, the refinement of FC metrics has practical implications for behavioral screening pipelines and genotype–phenotype mapping. Baseline-normalized indices, by reducing variance attributable to motor or anxiety-related traits, increase both sensitivity and reproducibility across experimental contexts. As demonstrated in our analyses, conventional freezing duration covaries strongly with locomotor activity rather than mnemonic processes (Fig. 2), underscoring the risk of interpreting raw freezing as a direct measure of learning. In contrast, the baseline-normalized indices isolate memory-related variance, allowing FC to more accurately capture cognitive performance. The observation that learning and memory align along a distinct behavioral dimension suggests that coarse-grained behavioral structure can, in principle, be recovered without exhaustive testing. However, this does not imply that individual phenotypes are fully characterized by a small number of assays. Indeed, within individual mouse strains, measures loading onto the same factor were not always consistent and in some cases showed opposing directions of effect. Thus, reduced test batteries may be sufficient for initial dimensional screening or broad phenotypic positioning, but detailed interpretation of genotype-specific phenotypes still requires task-level analyses. Notably, diverse learning-related measures, including spatial learning, avoidance learning, and motor learning tasks such as the rotarod, converged onto a single factor in our analysis. This convergence does not imply that these forms of learning are mechanistically identical, but rather that they share a common axis of behavioral variance at the phenotypic level. Accordingly, the extracted dimensions should be interpreted as organizing principles of behavioral expression rather than as direct proxies for discrete cognitive faculties.

### Limitations and future directions

Several limitations of this study should be noted. First, our analyses are based on strain-level summary statistics, which necessarily compress individual variability and learning dynamics into aggregate measures. As a result, within-strain heterogeneity and trial-by-trial learning trajectories are not directly captured. Future studies incorporating trial-level analysis, continuous behavior tracking, or time-resolved modeling approaches may provide deeper insight into the dynamics of learning and motor inhibition. Second, although baseline normalization reduces the influence of general locomotor activity, other factors such as nociceptive sensitivity, motivation, and stress reactivity may still contribute to freezing behavior. Combining refined FC metrics with physiological or neural measures, including calcium imaging or optogenetic manipulation, will be important for further disentangling these components. Finally, while the present framework emphasizes locomotor and cognitive domains, our dataset also includes social and affective assays. The fact that these domains did not emerge as primary axes suggests that resolving social or affective dimensions may require more specialized tasks or analytical approaches, rather than indicating their absence from mouse behavioral organization.

## Conclusion

Our findings challenge the widespread assumption that freezing duration is a direct and reliable indicator of fear memory. When interpreted using conventional freezing measures, fear conditioning does not provide a process-pure readout of memory but instead reflects a composite behavioral output shaped by locomotor, affective, and cognitive factors. By re-evaluating fear conditioning within a large-scale, multivariate framework, this study clarifies how learning-and memory-related performance and locomotor activity jointly structure mouse behavior. The results demonstrate that conventional freezing conflates motor and memory components, whereas baseline-normalized indices isolate memory-related variance and consistently align with a reproducible cognitive dimension. Together, these findings establish a scalable and conceptually grounded approach for interpreting complex behavioral datasets. Beyond refining the fear-conditioning paradigm, this dimensional framework provides a foundation for integrating behavioral, genetic, and neural data to advance a more mechanistic and reproducible behavioral neuroscience.

## Materials and Methods

### Animals and data collection

A total of 166 experimental mouse groups were analyzed (Table S2), including genetic mutants, drug-treated, psychostimulant-treated, and aged mice. Animals were housed under a 12 h light/dark cycle (7:00 AM to 7:00 PM) with ad libitum access to food and water. Adult male mice were used in all behavioral tests to eliminate the behavioral effects of the estrus cycle. Behavioral testing was performed under standardized environmental conditions, with test-specific parameters such as illumination, temperature, and humidity controlled as appropriate for each assay. Following general health and neurological screens (GHNS), mice were generally subjected to behavioral assessments in the following order: light/dark transition (LD), open field (OF), elevated plus maze (EP), hot plate (HP), social interaction in a novel environment (SI), rotarod (RR), Crawley’s 3-chamber social interaction test (CSI), startle response/prepulse inhibition (PPI), Porsolt forced swim (PS), T-maze (TM), Barnes maze (BM), tail suspension (TS), fear conditioning (FC), and home cage social interaction tests (HC), although slight modifications to the sequence or procedures were made in some strains or experiments. Measures obtained in each test are summarized in Table S1.

### Behavioral tests

#### General health and neurological screens (GHNS)

The presence of whiskers and bald patches was checked during GHNS and additionally when necessary as part of general health monitoring ^54^. The righting, whisker touch, and ear twitch reflexes were also evaluated as part of the neurological screening. Body weight and rectal temperature were measured. In addition, the neuromuscular function was assessed by grip strength and wire hanging tests. Briefly, grip strength was measured using a grip strength meter (O’Hara & Co., Tokyo, Japan). The mouse was positioned to spontaneously grasp a wire grid by the forelimbs and then pulled backward by the tail until wire release. The peak force was recorded in Newtons (N). Each mouse was tested three times, and the greatest value obtained was used for further analyses. In the wire hanging test, a mouse was placed on a wire mesh at the top of the apparatus (O’Hara & Co.), and the wire mesh was then gently turned upside down. The mouse gripped the wire in order not to fall off, and the latency to fall was recorded with a 60 s cut-off time.

#### Light/dark transition test (LD)

The light/dark transition test was conducted to measure anxiety levels as previously described ^55^. The apparatus consisted of two equal-sized plastic boxes (20 × 20 cm), one illuminated at approximately 390 lx and the other dark (< 2 lx), separated by a central partition plate with a small 3 × 5 cm opening allowing the mouse to transit from one box to the other. A mouse was first placed in the dark box and allowed to freely explore the apparatus for 10 min. The time to first entry into the light box, time spent in the light box, number of transitions between boxes, and distances traveled in light and dark boxes were measured using ImageLD software (see “Image analysis of behavioral tests”).

#### Open field test (OF)

Locomotor activity and exploratory behavior were measured using an open field apparatus (40 × 40 × 30 cm; Accuscan Instruments, Columbus, OH) as described previously ^56^. The test chamber was illuminated at approximately 100 lx. A mouse was placed in the corner of the apparatus and total distance traveled (cm), time spent in the center area (20 × 20 cm), vertical activity, and stereotypic counts over 120 min were recorded by the VersaMax.

#### Elevated plus maze test (EP)

An elevated plus maze test was conducted as previously described ^57^. The apparatus consisted of two open arms (25 cm long × 5 cm wide) crossing two enclosed arms of identical dimensions but with 15-cm high transparent walls, all connected by a central platform. The entire apparatus was elevated 50 cm above the floor, and the illumination at the central platform was set to approximately 100 lx ^58^. A mouse was placed at the center of the maze, facing one of the closed arms, and allowed to freely explore for 10 min. The numbers of open-and closed-arm entries, distance traveled, percentage of open-arm entries, and percentage of time spent in the open arms were measured using ImageEP software (see “Image analysis of behavioral tests”).

#### Hot plate test (HP)

Pain sensitivity was measured by the hot plate test. A mouse was placed on a 55°C hot plate (Columbus Instruments, Columbus, OH), and the latency to the first nociceptive response was recorded with a 15-s cut-off time.

#### Social interaction test in a novel environment (SI)

The one-chamber social interaction test in a novel environment was performed as described previously ^54^. Two mice of the same condition (e.g., genotype) but reared in different cages were placed together in a box (40 × 40 × 30 cm) and allowed to explore freely for 10 min. Mouse behaviors were recorded with a CCD camera. The total duration of contact, the total number of contacts, the total duration of active contact, the mean duration per contact, and the total distance traveled were measured automatically using ImageSI software (see “Image analysis of behavioral tests”). Active contact was defined as maintained contact while either mouse traveled more than 10 cm between two successive image frames acquired 3 frames per second.

#### Rotarod test (RR)

Motor coordination and motor learning were examined using an accelerating rotarod (UGO Basile, Comerio, VA, Italy). The latency to fall from the rod was recorded during three daily trials conducted on 2 consecutive days (3 × 2 trials). During each trial, rod speed was increased from 4 to 40 rpm over the 5-min test period.

#### Crawley’s three-chamber social interaction test (CSI)

Crawley’s three-chamber social interaction test was performed to assess sociability and preference for social novelty as previously described ^56,59,60^. The experimental apparatus was a 41 × 62 cm rectangular non-transparent gray Plexiglas box separated into three equal-sized chambers (20 × 40 cm) by transparent Plexiglas plates with small openings allowing mice to freely transit from one chamber to another. One wire cylinder-shaped cage was placed in the corner of each side chamber. Tested mice were first allowed to freely explore the chambers for 10 min as a habituation period. A stranger mouse was then placed randomly in one of the wire cages in a side chamber, and the tested mouse was again allowed to freely explore the chambers for 10 min. In the final session, a second stranger mouse was placed in the wire cage in the opposite side chamber, and the tested mouse was again allowed to freely explore the chambers for 10 min. The numbers of contacts and the time spent interacting with strangers 1 and 2, as well as the distance traveled in each chamber, were recorded using ImageCSI software (see “Image analysis of behavioral tests”).

#### Startle response / prepulse inhibition test (PPI)

Acoustic startle response and prepulse inhibition were assessed as previously described^54^. Prepulse inhibition measures the reduction in the startle response when a weak prepulse precedes a louder startle stimulus. The system used for the detection of startle reflexes (O’Hara & Co.) consisted of a standard cage mounted on a movement sensor in a sound-attenuated chamber with fan ventilation. A test session was composed of six trial types: two startle stimulus-only trials (110 or 120 dB) and four prepulse inhibition trials (74 or 78 dB prepulses delivered prior to a 110-or 120-dB startle). Six blocks of the six trial types (i.e., 36 trials in total) were conducted, with each trial type presented once in pseudorandom order within each block. The PPI (%) was calculated for each trial type according to the following equation: {[(startle amplitude of trial without prepulse) – (startle amplitude of trial with prepulse)]/(startle amplitude of trial without prepulse)} × 100.

#### Porsolt forced swim test (PS)

The Porsolt swim test was performed as previously described ^54^. The apparatus consisted of four Plexiglas cylinders (12 cm diameter × 22 cm height) filled with water at approximately 23°C to a depth of 7.5 cm. Four mice were placed individually in the cylinders, and images were captured at two frames per second for 10 min. Swimming distance and immobility time (% of total) were recorded automatically using ImageTS software (see “Image analysis of behavioral tests”).

#### T-maze forced alternation / spontaneous alternation test (TM)

A T-maze forced or spontaneous alternation task was performed as previously described ^61^. The T maze apparatus consisted of three white plastic runways with 25-cm high walls (O’Hara & Co.), partitioned into six areas by sliding doors. The stem of the T comprised a start compartment (S1) at the arm intersection and a main stem area (13 × 24 cm; S2). The two arms similarly comprised passage areas (P1 and P2) adjacent to S1 and main areas (A1 and A2; 11 × 20.5 cm). In each trial, the mouse was allowed to choose between the two goal arms, and entry into the arm opposite to that visited in the preceding run was scored as an alternation response, whereas re-entry into the same arm was scored as an error. Repeated sessions were conducted with or without inter-trial delays depending on the strain and experimental design. Data were acquired and sliding doors were controlled automatically using ImageTM software (see “Image analysis of behavioral tests”).

#### Radial maze test (RM)

Working memory was assessed in some strains using a radial arm maze test rather than a T-maze test, depending on the strain and experimental cohort. The test was conducted as described elsewhere ^62^. For the present analyses, latency to obtain all pellets, total distance traveled, and the number of working memory errors were used as measures corresponding to latency, total distance traveled, and the number of errors in the T-maze. Details of which assay was used for each strain are provided in Table S2.

#### Barnes maze test (BM)

Spatial learning and memory were assessed using the Barnes maze as described previously ^56^. The maze was a circular whiteboard (1m in diameter) with 12 holes equally spaced around the margin and elevated 75 cm from the floor (O’Hara & Co.). A black plastic box (17 × 13 × 7 cm) lined with paper cage bedding was positioned under one of the holes (the target). The target hole (1–12) was fixed for each mouse but assigned pseudo-randomly across individuals. The apparatus was brightly illuminated (approximately 800 lx or higher). The board was rotated daily so that the spatial location of the target changed relative to distal visual room cues while proximal cues were held constant. Training schedules, including the number of trials per day and the total number of training days, varied depending on the strain and experimental cohort. Each trial lasted a maximum of 5 min and was completed when the mouse entered the black plastic box. Probe tests were conducted for 3 min without the black plastic box one day or one month after the final training trial. The number of errors, latency, and distance traveled to reach the target hole for the first time, the number of omission errors and time spent around each hole were recorded by ImageBM software (see “Image analysis of behavioral tests”).

#### Tail suspension (TS)

Mice were suspended 30 cm above the floor in a visually isolated area by adhesive tape placed approximately 1 cm from the base of the tail. Behavior during suspension was recorded for 10 min using ImageTS software (see “Image analysis of behavioral tests”) and immobility time (% of total time) was measured automatically.

#### Fear conditioning test (FC)

Contextual and cued fear conditioning tests were conducted and analyzed as described in a previous study ^54,63^. Mice were placed in a transparent acrylic chamber (33 × 25 × 28 cm, O’Hara & Co.) with background white noise at 55 dB and illumination at 100 lx. In the conditioning session, the conditioned acoustic stimulus (CS, 55 dB) was presented for 30 s at three times (2, 4, and 6 min), and a mild foot shock (2 s, 0.3 mA) was presented as the unconditioned stimulus (US) at the end of each CS. One day or a month after the conditioning session, contextual fear was measured in the same chamber, while cued fear was measured thereafter in a triangular box (33 × 29 × 32 cm) constructed of white opaque Plexiglas under approximately 30 lx illumination. Mice were placed in the chamber for 3 min with neither CS nor US presented. Thereafter, the CS (55 dB) was presented for the last 3 min and images were captured at 1 frame per second. Each pair of successive frames in which the mouse moved was measured. When this measure was below a certain threshold (20 or 30 pixels), the mouse was considered “freezing”; alternatively, when this measure equaled or exceeded the threshold, the behavior was considered “non-freezing”. “Freezing” that lasted less than 2 s was not included in the analysis. Data acquisition, control of stimuli (i.e., tones and shocks), and data analysis were conducted automatically using ImageFZ software (see “Image analysis of behavioral tests”).

#### Home cage social interaction test (HC)

The 24-hour home cage locomotor activity and social interaction test was conducted as previously described ^64^ for one continuous week. The monitoring system included a standard home cage (29 × 19 × 13 cm) with a filtered cage top and an infrared video camera (O’Hara & Co.). Two mice of the same condition (e.g., genotype) that had been housed separately were placed together in the home cage, and locomotor activity and social behavior were assessed by automated image analysis. Social interaction was quantified based on the number of particles detected in each frame: two particles indicated that the mice were apart, whereas one particle indicated physical contact, because the two mice were detected as a single particle when in contact. Data acquisition and analysis were conducted using ImageHA software (see “Image analysis of behavioral tests”)

#### Image analysis of behavioral tests

In-house programs were used to analyze the imaging data acquired for several behavioral tests (ImageLD, ImageEP, ImageSI, ImageCSI, ImageTS, ImageTM, ImageBM, ImageFZ, and ImageHA) based on ImageJ (U.S. National Institutes of Health; available at https://imagej.nih.gov/ij/), modified by Tsuyoshi Miyakawa (available through O’Hara & Co.).

#### Effect sizes calculated for behavioral measures

Sixty-eight behavioral indices from 15 behavioral tests were included in the analysis (see Table S1). For each behavioral measure, the difference between an experimental group and its corresponding control group was quantified as Hedges’ adjusted *g*, a less biased estimator of effect size ^65^. When the same experimental group was assessed in multiple independent batches, batch-specific effect sizes were combined into a pooled effect size by random-effects meta-analysis using the metafor package ^66^ in R. Experimental groups with data from at least six behavioral tests were included, yielding 166 experimental group-control comparisons for subsequent analyses. Details of the experimental groups and corresponding controls are provided in Table S2.

#### Systematic literature search and categorical classification of fear conditioning studies

We systematically searched the Europe PubMed Central (PMC) database for full-text articles published between 2020 and 2025. The search targeted experimental studies that included fear conditioning assays. Articles were retrieved in XML format through the Europe PMC RESTful API for automated processing. A custom Python-based pipeline was used to extract relevant text and classify studies according to how locomotor activity was assessed and incorporated into the interpretation of fear conditioning measures, as described below. The retrieved studies were initially classified into three categories.

Category 1: Use of locomotor-adjusted fear conditioning indices (e.g., activity suppression ratios or baseline-normalized freezing).

Category 2: Independent assessment of locomotor activity (e.g., Open Field Test, distance traveled), further subdivided according to its analytical relationship to fear conditioning measures:

− 2a: locomotor activity included as a statistical covariate in the analysis of fear conditioning (e.g., ANCOVA);
− 2b: significant group differences in locomotor activity reported;
− 2c: no significant group differences in locomotor activity reported;
− 2d: locomotor activity mentioned or measured, but not explicitly related to fear conditioning outcomes in the statistical analysis.

Category 3: No mention of locomotor assessment in relation to freezing behavior. Because automated text mining can over-detect terms such as activity suppression in non-FC contexts, all candidate Category 1 and 2a studies were manually reviewed before final counting. This review was used to confirm that the studies indeed employed locomotor-adjusted fear conditioning indices or incorporated locomotor activity as a statistical covariate in the analysis of fear conditioning outcomes.

### Multiple factor analysis (MFA) and hierarchical clustering for behavioral measures

To quantify the dis-/similarity of behavioral patterns among mouse strains, we first performed factor analysis based on the effect sizes between control and mutant mice. As a preliminary step, the imputation of missing values was conducted with the imputeMFA function in the missMDA package (version 1.18) in R. We then performed multiple factor analysis (MFA) ^67^ using the MFA function of the factoMineR (version 2.4) package in R, supposing that each set of behavioral measures constituting a test corresponds to variables structured in a group. The MFA is one of the factorial methods that generalize principal component analysis or multiple correspondence analysis. It aims to balance the influences of each set of variables measured in the same observation by normalizing them with weight. In our analysis, MFA was applied to 68 behavioral measures grouped into 15 tests, with the number of dimensions set to four. Of these, the first two dimensions were retained as representative behavioral components. The third and fourth dimensions were excluded from subsequent analyses because individual behavioral measures generally showed low contributions to and weak correlations with these dimensions, and no clear associations with the fear-conditioning measures were identified.

## Acknowledgement

We are grateful to Dr. Makoto Noda, Dr. Daisuke Yabe, Dr. Yasuo Tsutsumi, Dr. Hiroshi Hosokawa, Dr. Yoshihiro Yoshihara, and Dr. Motoi Okamoto for providing valuable mouse behavioral datasets.

## Contributions

Daiki X. Sato: Conceptualization, Methodology, Software, Formal Analysis, Data Curation, Visualization, Writing – Original Draft, and Writing – Review & Editing. Markos M. Chatzigiannis: Methodology, Formal Analysis, Data Curation, and Writing – Original Draft. Hirotaka Shoji: Investigation, Resources, Data Curation, and Writing – Review & Editing. Giovanni Sala: Conceptualization, Methodology, Software, Formal Analysis, and Data Curation. Satoko Hattori and Keizo Takao: Investigation, Resources, and Data Curation. Tatsuo Kinashi, Tatsuya Kishino, Sumiyo Morita, Izuho Hatada, Yo Shinoda, Kotaro Hattori, Takeshi Yagi, Akinobu Matsumoto, Hideo Egawa, Shoko Nishihara, Kimiko Shimizu, Koji Ikegami, Maki K. Yamada, Hiroshi Ageta, Mitsutoshi Setou, Hirotaka Tao, Naoto Ueno, Pradeep Bhandari, Ryuichi Shigemoto, Shuji Wakatsuki, Toshiyuki Araki, Akihiro Yamanaka, Hideyuki Mukai, Tadahiro Nagaoka, Masashi Kishi, Shigeki Furuya, Tomomi Yamamoto, Yoshihiro Kubo, Yuichi Iida, Yasuhiro Kazuki, Hideki Enomoto, Motoaki Fukasawa, Nobuteru Usuda, Satoshi Inoue, Kaoru Inokuchi, Tsuyoshi Hattori, Mariko Taniguchi-Ikeda, Tatsushi Toda, Akiko Kubo, Katsuhiro Kawaai, Katsuhiko Mikoshiba: Investigation and Resources. Louie N. van de Lagemaat, Noboru H. Komiyama, and Seth G. N. Grant: Resources, Data Curation, and Writing – Review & Editing. Tsuyoshi Miyakawa: Conceptualization, Resources, Writing – Review & Editing, Supervision, Project Administration, and Funding Acquisition. All authors reviewed and approved the manuscript.

## Funding

This work was supported by grants to T.M. from the Ministry of Education, Culture, Sports, Science and Technology (MEXT) Promotion of Distinctive Joint Research Center Program (Grant Numbers JPMXP0618217663 and JPMXP0621467949); the Japan Society for the Promotion of Science (JSPS) KAKENHI (Grant Numbers JP16H06276, JP22H04922 [AdAMS], JP20H00522, JP25K00903, JP16H06462, JP15H01297, JP20016013, JP20023017, JP16680015, JP19653081, and JP21300121); the Japan Agency for Medical Research and Development (AMED) (Grant Number JP21dm0107101); Integrative Brain Research (IBR-shien); the Comprehensive Brain Science Network; the Promotion of Fundamental Studies in Health Sciences of the National Institute of Biomedical Innovation (NIBIO); the Neuroinformatics Japan Center (NIJC); and CREST and BIRD of the Japan Science and Technology Agency (JST). This work was also supported by JSPS KAKENHI Grant Numbers JP22116511 to N.Ue., JP18022025 to H.M., JP25430033 to M.K., and JP18300125 and JP24K01703 to S.F.; the Austrian Science Fund (FWF) Grant Number PAT5720324 to R.S.; and AMED Grant Number JP26ama121046 to Y.Ka. The G2C Program (NHK, SGNG) received funding from The Wellcome Trust and the EU FP7 Framework Programmes EUROSPIN (FP7-HEALTH-241498), SynSys (FP7-HEALTH-242167), and GENCODYS (FP7-HEALTH-241995).

## Competing interests

The authors declare no competing interests.

## Data and code availability

Data and codes to reproduce the findings described in the present study will be available upon publication.

---

**Figure S1.**
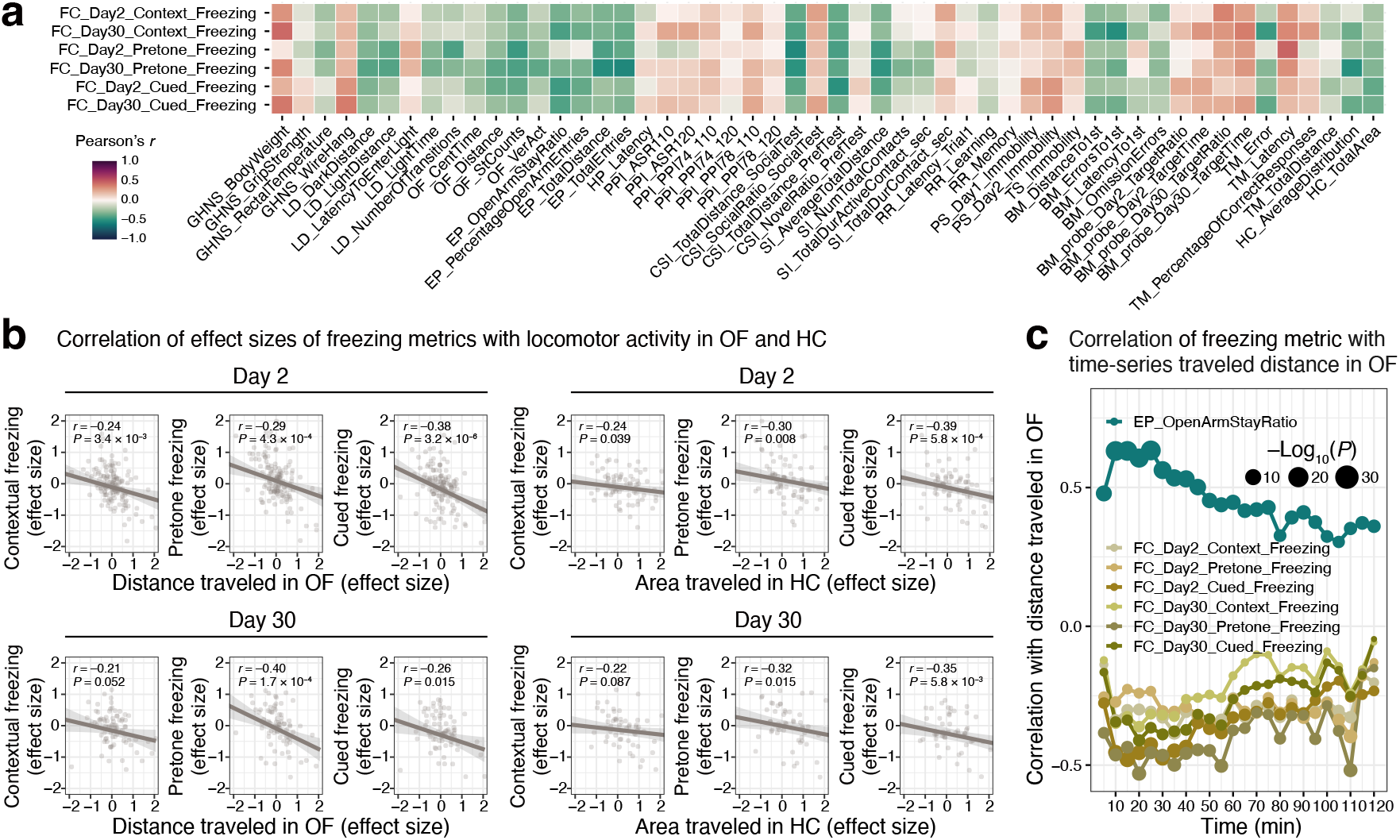
Conventional freezing metrics covary with locomotor and exploratory measures across behavioral tests. **(a)** Heat map showing Pearson correlations between strain-level effect sizes (Hedges’ g) of fear-conditioning (FC) freezing measures (context, pretone, and cued freezing on Day 2 and Day 30) and effect sizes of behavioral variables from the other tests in the battery. Colors indicate Pearson’s *r*. **(b)** Scatterplots showing correlations between effect sizes of FC freezing measures and representative locomotor indices from open field (OF_Distance) and home cage (HC_TotalArea). Each point represents a mouse strain; *r* and P values are shown in each panel. **(c)** Time-resolved correlations between OF traveled-distance effect sizes calculated in 5-min bins across the OF session and effect sizes for FC freezing and EP open-arm duration. Lines show Pearson’s *r* as a function of time. Point size indicates statistical significance (–Log10(*P*)).

**Figure S2.**
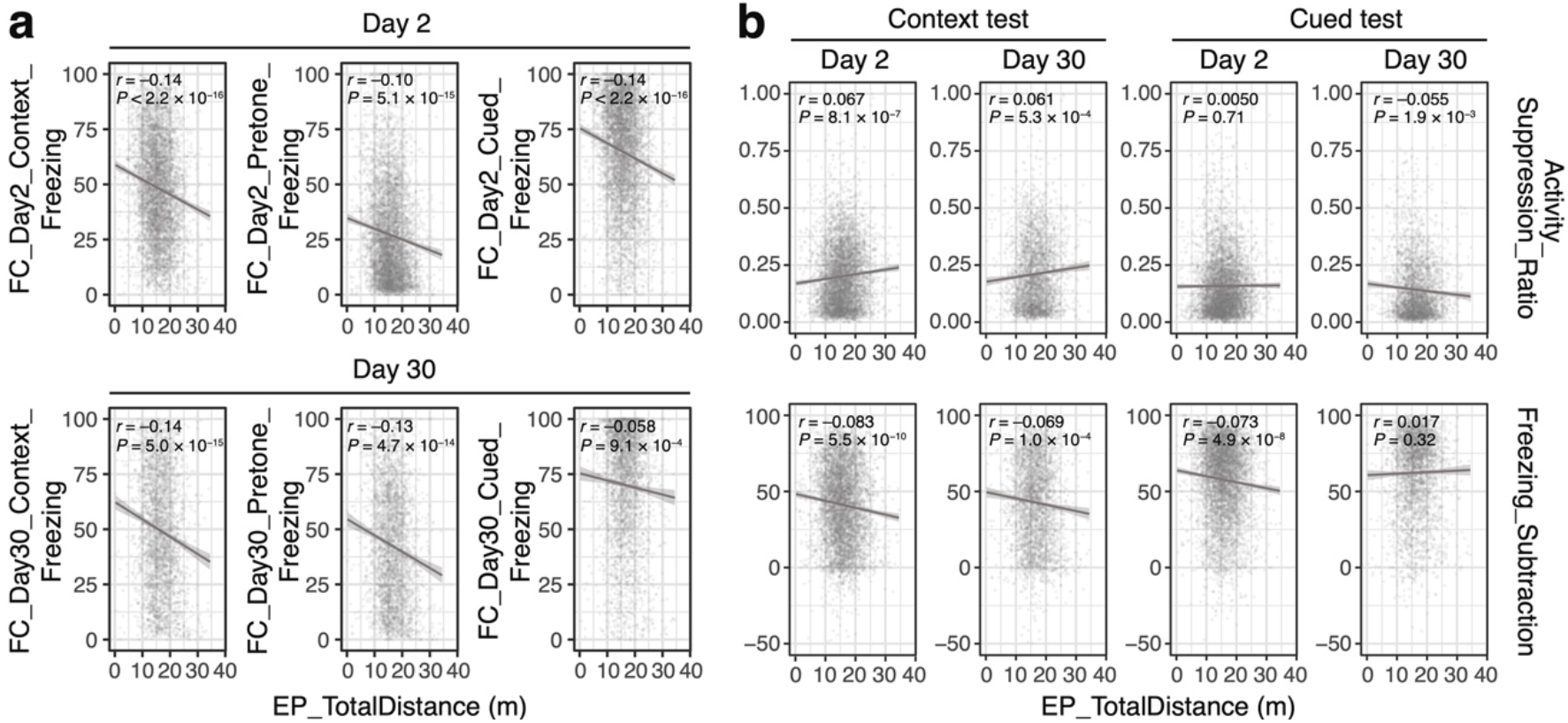
Individual-level correlations show that baseline-normalized fear-conditioning metrics minimize locomotor confounds. Scatterplots of individual-level data showing correlations between freezing-related measures and total distance traveled in the elevated plus maze (EP_TotalDistance). **(a)** Conventional freezing scores showed strong negative correlations with activity at both Day 2 and Day 30, while **(b)** baseline-normalized indices (Activity Suppression Ratio and Freezing Subtraction) showed little or no dependence on activity, consistent with the strain-level results in Fig. 2.

**Figure S3.**
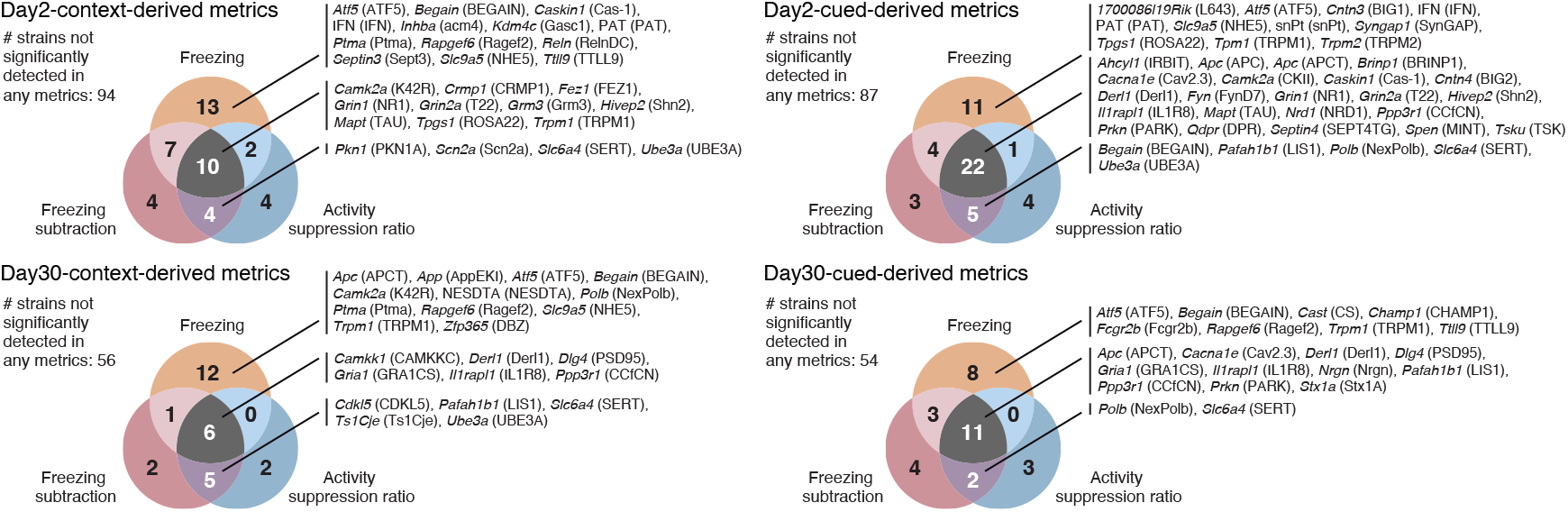
Baseline-normalized fear conditioning metrics detect synaptic plasticity mutants involved in learning and memory. Venn diagrams showing overlap among strains with significant genotype effects across conventional freezing, Freezing Subtraction, and Activity Suppression Ratio for context and cued tests at Day 2 and Day 30. Each panel lists the genes related to the mutant strains identified in each category. Baseline-normalized and conventional metrics commonly detected classical plasticity-related mutants such as *Grin1*, *Grin2a*, *Camk2a*, *Dlg4*, *Gria1*, and *Ppp3r1*, indicating improved specificity for memory-related phenotypes compared with conventional freezing scores.

## Notes

### Competing Interest Statement

The authors have declared no competing interest.

