## Supplementary figures and images for "Large-scale dimensional behavioral profiling dissociates fear memory from locomotor confounds in mice: The necessity of baseline-normalized metrics"

### Supplementary Figure S1

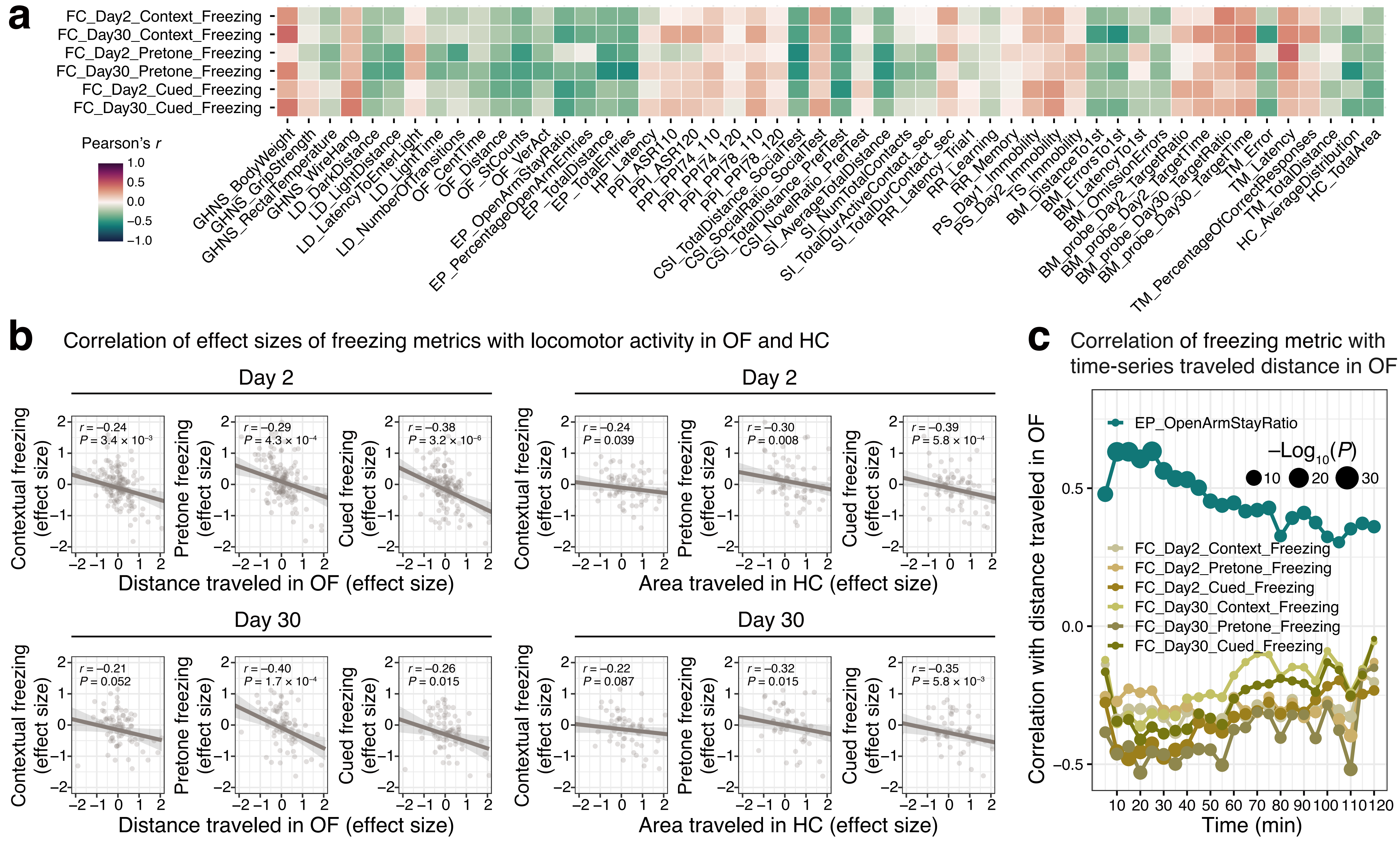

### Supplementary Figure S2

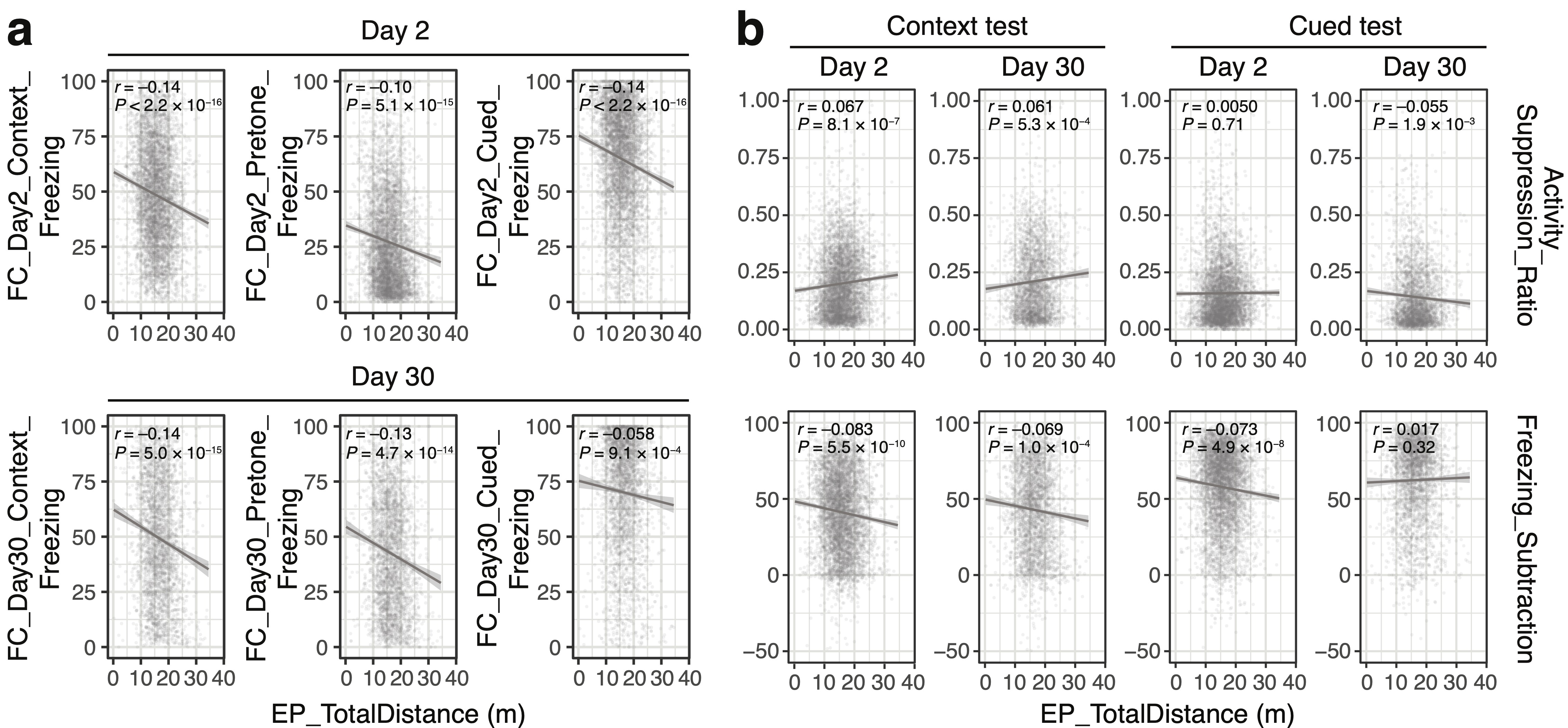

### Supplementary Figure S3

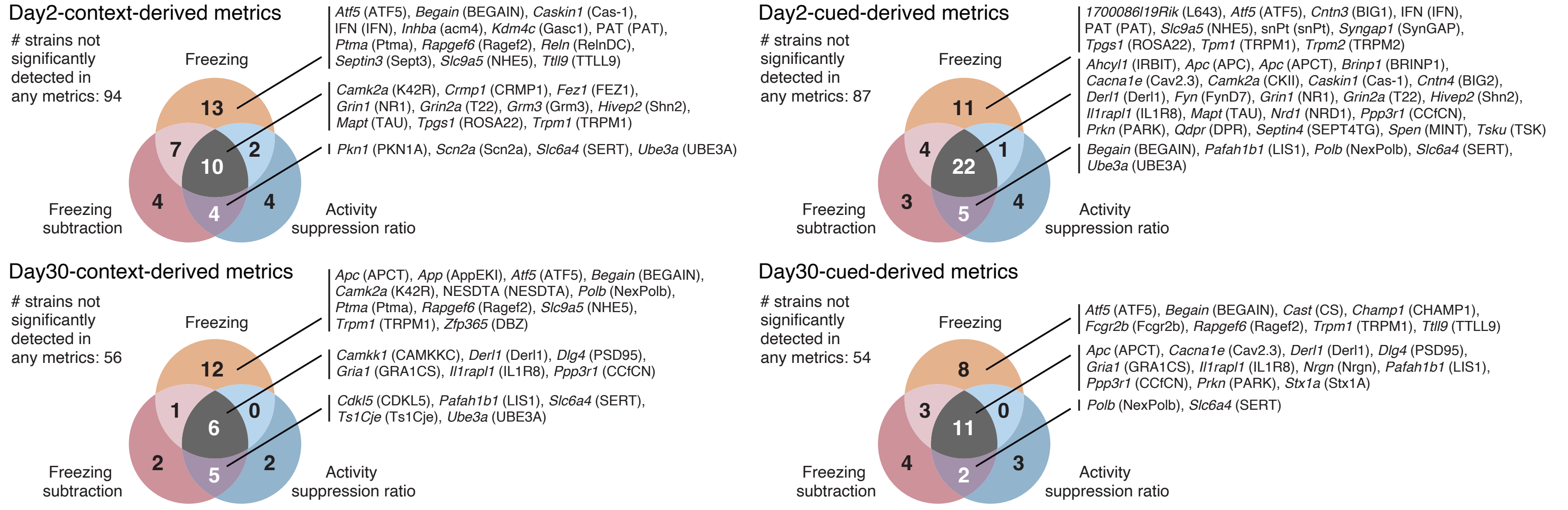
